# Poor hive thermoregulation produces an Allee effect and leads to colony collapse

**DOI:** 10.1101/2020.01.21.914697

**Authors:** Zeaiter Zeaiter, Mary R. Myerscough

## Abstract

In recent years the honey bee industry has bee experiencing increased loss of hives. The accumulation of multiple stressors on a hive potentially drives hive loss in various ways, including winter loss and colony collapse disorder. One of these stressors is the breakdown of thermoregulation inside the hive. For pupae to develop correctly into healthy adult bees, the temperature within the hive must be regulated by the hive bees to within a narrow range that ensures optimal development. Suboptimal development in adults affects their brain and flight muscles so bees becomes inefficient foragers with shorter life spans. We model the effect of thermoregulation on hive health using a system of delay differential equations that gives insights into how varying hive temperatures have an effect on the survival of the colony. We show that thermoregulatory stress has the capacity to drive colony loss in the model via a saddle-node bifurcation with an associated Allee effect.

## 1. Introduction

There are increasing pressures on honey bees in today’s industrialised world. The loss of honey bees has a profound effect, not only on natural systems but also on food production. The Western honey bee, *Apis mellifera*, is the most common commercial pollinator [4]. Pollinators are particularly vital in agriculture to grow commercial quantities of many crops, particularly fruit, vegetables and nuts.

In recent years honey bee colonies have been failing at alarming rates and countries have begun monitoring their honey bee stocks [18]. One conceptually important mode of colony loss is colony collapse disorder (CCD). Colony collapse disorder is characterised by a vacant hive with dead brood and stored food present, but few to no adult bees, which suggests a rapid depopulation [1]. In some cases a colony will fail so rapidly that within the space of a few weeks an apparently healthy hive becomes emptied of adult bees [1, 2, 3].

Mathematical models that represent the dynamics of a honey bee colony are a useful tool to examine causes of hive loss with a focus on CCD. Khoury *et al.* [5, 16] produced two foundational models. The first of these, models populations of adult hive bees and foragers. The second also includes the brood population and stored food. These models have been extended in several different ways to include seasonality [9], disease and infection [9], stressors [7, 8], and age-dependent foraging [6].

It is now thought that CCD is caused by the accumulation of stressors on the hive, rather than having one single cause. Booton *et al.* [8] and Bryden *et al.* [7] both model the effects of stress on a hive, but neither model has an explicit stressor. Booton *et al.* use a per capita death rate due to a generalised, unspecified stressor. Bryden *et al.* split the adult population into a healthy and impaired class based on a constant impairment rate.

Honey bee colonies have been very well studied over time and so many of the stressors that a hive experiences are well known and their effects, increasingly, are being quantified [20, 21]. Consequently, specific known hive stressors can be included directly in models, and, we would argue, that this approach will result in more useful models than a generic stress whose source is unknown. In this paper we will model, explicitly, one of these known stressors, the breakdown of thermoregulation, and explore its effects on the hive.

### 1.1. Understanding Thermoregulation and its Effect

Honey bees maintain precise environmental conditions inside the hive including, humidity, temperature and carbon dioxide levels [11]. Thermoregulation has a significant impact at particular points in the honey bee life cycle. The queen bee lays eggs which hatch into larvae (usually referred to as uncapped brood). The larvae then pupate. The brood cell containing the larvae is sealed by hive bees. Hence this pupal stage is referred to as capped brood. After 12 days an adult bee emerges from the capped cell. The newly emerged adult bee will initially work within the hive and later is recruited to foraging duties.

Hive thermoregulation by workers in general, allows the hive to survive colder temperatures. However, a more specific role is to provide the optimal temperature for capped brood to develop into adult bees. For optimal development, capped brood must be maintained between 34 °C and 36 °C [10]. The process of thermoregulation requires more food consumption as the hive bees’ metabolic rate increases [10, 12] to meet the energetic requirements of hive thermoregulation.

A breakdown in thermoregulation has deleterious effects on the adult bees which emerge from the capped brood cells. Adult bees that have not developed at optimum temperatures show decreased cognitive ability, reduced foraging capability and higher susceptibility to parasites [11, 13, 14]. Capped brood that develop at lower than ideal temperatures will be inferior foragers later in life, collect less food per forager and have a shorter lifespan than their optimally developed counterparts. Jones *et al.* [11] conducted a study on the short-term and long-term memory of adult worker bees who experienced temperatures of 31, 32, 33, 34, 35, 36 and 37 °C*±*0.5 °C, respectively, during their time as capped brood. The results showed that there was no significant difference in long-term memory, but that there was a significant difference between the short-term memory of brood reared at 32 °C, 33 °C and 36 °C, with the latter having the best short-term memory of the three cohorts.

When foragers that are seeking new sources of pollen and nectar for the hive, find a suitable source they return to the hive and perform a dance to communicate what they have found. If successful, the forager will recruit other foragers to forage the new source of pollen and nectar. Tautz *et al.* [14] examined the effect of different rearing temperatures on brood and their ability, as foragers, to effectively communicate via dance. The authors showed that foragers who were raised at 32 °C had approximately 60% probability of performing a dance after visiting a feeder as opposed to the two other cohorts (one reared at 36 °C and the other reared under natural conditions) who displayed a 90% probability of performing a dance. Further, there were significantly smaller mean numbers of dance circuits by bees reared at 32 °C compared to those reared at 36 °C. This suggests that information about food sources will not spread as well in hives whose thermoregulation is sub-optimal.

The prevalence of a parasitic infection may also be attributed to poor thermoregulation. McMullan *et al.* [13] showed that a change in brood temperatures from 34 °C to 30 °C saw an increase in the prevalence of tracheal mites (20% to 41%, respectively).

It is evident from these studies that poor thermoregulation of capped brood impacts hive dynamics, including food collection, when these pupae become adults. Further it is clear that poor thermoregulation puts the hive under stress in particular and known ways.

There have been many mathematical models that capture the dynamics of a hive. Khoury *et al.* [5, 16] presented two models which modelled population dynamics with and without food in the hive. Both models made claims about the viability of a hive as a function of forager mortality. Perry *et al.* [6] extended the models proposed by Khoury *et al.* by including, explicitly, the role of worker age at onset of foraging. This was done by introducing three age-dependent functions. The idea of average mortality, which gives a lifespan of 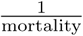, was used to define age at onset of foraging. If we consider the rate of recruitment of hive bees to become foragers (effectively the “mortality” of hive bees), then it follows that the average time that a bee spent as a hive bee is the average age at onset of foraging.

In this paper we will incorporate these known outcomes of thermoregulatory stress into a mathematical model. We aim to extend the work presented in previous models [5, 6, 16] by introducing the effects of a breakdown of thermoregulation within the hive. We use the model presented by Perry *et al.* [6] as a foundation and incorporate thermorelatory stress explicitly.

## 2. Methods

We will use the model of Perry *et al.* [6] as a foundation for our model. This model has four dependent variables: food, uncapped brood, hive bees and foragers (denoted by the variables *f*, *B*, *H* and *F*, respectively). The model explicitly includes the age at onset of foraging of hive bees. This was derived using the average rate, *R*(*H, F, f*), that hive bees leave that class. This is the rate of recruitment to foraging,

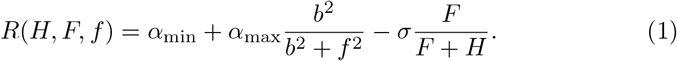

The first and second term, respectively, represent the standard rate of recruitment and the accelerated rate when there is a low amount of stored food in the hive. The last term represents the effect of social inhibition on forager recruitment. Therefore the age at onset of foraging is defined as,

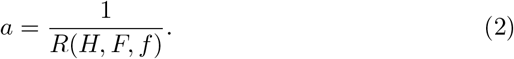

The equation for the rate of change of stored food is,

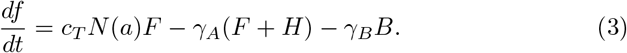

The first term on the right hand side models the rate of food collection, where *c*_*T*_ is the amount of food collected per forager per trip and *N*(*a*) is the number of trips a forager makes per day as a function of age. The last two terms on the right hand side are the rate of food consumption where, *γ*_*A*_ and *γ*_*B*_ is the amount of food consumed per day per adult bee and brood, respectively.

The equation for the rate of change of the uncapped brood population is defined as,

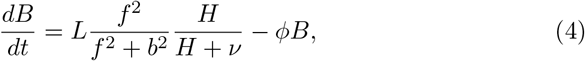

where first term represents the number of eggs laid per day, *L*. This is dependent on the availability of food and hive bees in the hive. The second term represents the rate that uncapped brood progress to capped brood at a pupation rate *φ*. The brood which pupate do not immediately join the hive bee class because capped brood require approximately 12 days to mature into adult bees. The equation for the rate of change of the hive bee population is,

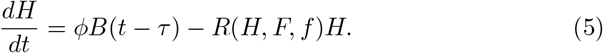

The first term on the right hand side represents the rate that uncapped brood entered pupation *τ* = 12 days ago and hence the rate that hive bees are currently emerging from pupation. The second term is the rate the hive bees progress to foraging duties at the recruitment rate *R*(*H, F, f*).

Finally the the equation for the rate of change in the population of foragers is,

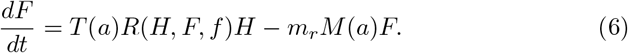

The first term on the right hand side is the rate that hive bees are recruited to foraging duties following their orientation flights. The function *T* (*a*) represents the transitional survival rate as a function of age of onset of foraging, *a*. The last term represents the death rate of foragers, where *M*(*a*) is the mortality rate as a function of *a* and *m*_*r*_ is the ratio of the death rate of a stressed hive to the death rate of a healthy hive. We now identify where thermoregulatory stress acts in the model and construct our stress function.

### 2.1. Incorporating Stress

We aim to extend the model presented by Perry *et al.* by introducing a function which models the effects of a breakdown in thermoregulation. We can identify three components of the model which are affected by thermoregulatory stress. Let us denote three stress functions *S*_1_, *S*_2_ and *S*_3_. We will specify each of these functions in detail later.

Studies have shown that poor thermoregulation leads to inferior foragers with lower cognitive ability [11, 14]. These inferior foragers may choose poorer pollen and nectar sources and not be able to forage as efficiently as their optimally developed coutnerparts. Hence, we assume that the food collection term in (3) is inversely proportional to the stress function *S*_1_,

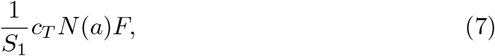

so that when the impact of thermoregulatory stress is high, the inferior foragers will collect less food per forager.

For the hive to rear properly developed brood it must be maintained at temperatures that require active thermoregulation. This creates a need to actively warm the hive and so hive bees will regulate the hive temperature by metabolising sugars rapidly to generate heat with their flight muscles. This leads to an increase in the caloric needs of the hive bees and higher food consumption. Assuming that there is a linear relationship between the change in hive temperature and the food consumption of hive bees, we multiply the food consumption of hive bees by the stress function *S*_2_,

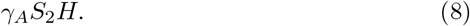

Finally, we assume that the foragers that have pupated under a regime of poor thermoregulation have a reduced lifespan, and so we multiply their mortality term by the stress function *S*_3_,

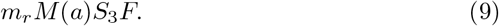

We have outlined the components of the Perry *et al.* model where we know that thermoregulatory stress acts, and how we should model this. However, we need to identify in detail how the effect of this stress should be modelled. In particular, we will need two functions. One which models the effect of stress due to the behaviour of the current hive bee population and one which is dependent on historical hive bee populations.

### 2.2. Breakdown in Thermoregulation

As the number of hive bees within a hive diminishes, the efficacy of thermoregulation also diminishes. This causes an increase in the effects of stress on the hive. We assume that there are bounds on the maximum and minimum effects of stress. We aim to model stress using an inverse sigmoid function of the form,

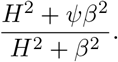

As identified earlier, the three areas of impact of thermoregulatory stress in the model are food collection, food consumption and forager mortality. These three areas are not all effected by stress in the same way. If we assume a hive is experiencing a breakdown in thermoregulation then the current hive bee population will be actively regulating temperature. The current forager population may also have experienced some form of stunted development as a result of a prior hive bee cohort. So the current hive bee population will have an increased food consumption as a result of thermoregulating and the current forager cohort may be sub-optimally developed and inefficient foragers. Therefore, we seek to derive two different stress functions, one dependent on the current hive bee population, and the other dependent on historical hive bee populations.

We first look at how past thermoregulation history affects current adult bee populations. In particular, to capture the effects of poor thermoregulation on foragers, we need to have information about the hive bee population that was present when the current foragers were capped brood.

Figure 2 shows this relationship. We will use *τ* exclusively to represent delays. Let *τ*_*y*_ and *τ*_*o*_ (respectively) denote the amount of time since the youngest and oldest of the current foragers entered pupation. In order to represent the hive bee population when foragers were capped brood we use distributed delays.

**Figure 1:**
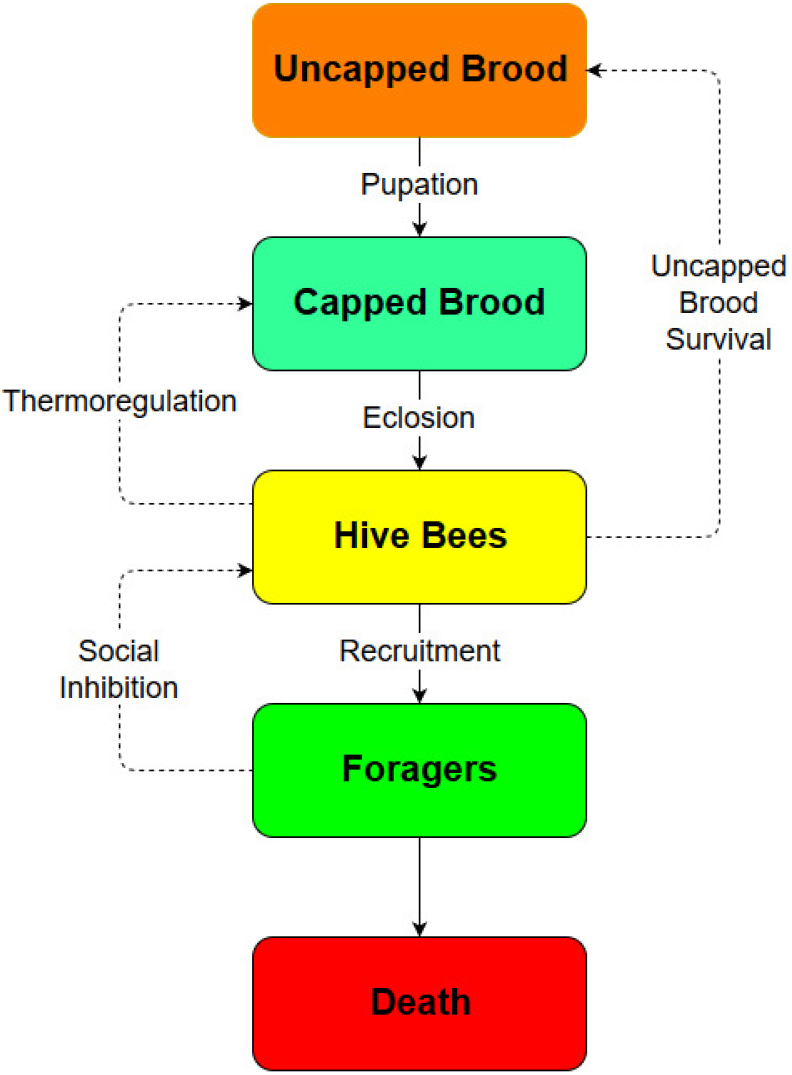
Life cycle of model honey bee. We see where thermoregulation acts. Hive bees thermoregulate to maintain an optimal temperature to allow the capped brood to develop properly.

**Figure 2:**
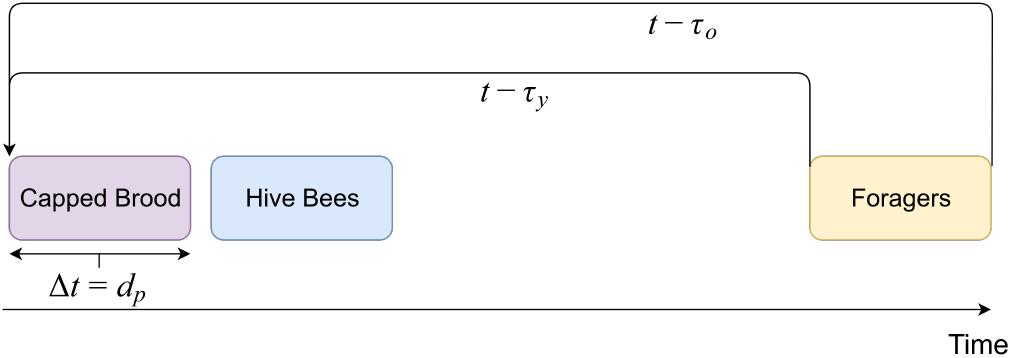
We track the hive bee population at the time the current forager cohort were capped brood. The youngest of the current foragers were brood entering pupation *t* − *τ*_*y*_ days ago, and the oldest of the current foragers were brood entering pupation *t* − *τ*_*o*_ days ago.

The first distribution in delay arises because the forager cohort contains bees of different ages. The second distributed delay is generated by the duration of pupation for each age group of current foragers.

To formulate the first of these distributed delays we look at the case of the youngest and oldest foragers. The youngest foragers entered pupation *τ*_*y*_ days ago, and so there were *H*(*t* − *τ*_*y*_) hive bees present when they entered pupation. Letting *d*_*p*_ denote the duration of pupation, then there would have been *H*(*t* − (*τ*_*y*_ − *d*_*p*_)) hive bees present when the youngest current foragers finished pupation. So the average number of hive bees present for the entire duration of pupation of the youngest foragers is,

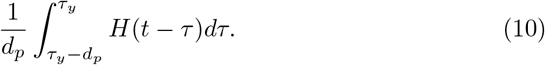

Likewise, for the oldest current foragers, the average number of hive bees present for the entire duration of pupation of the oldest current foragers is,

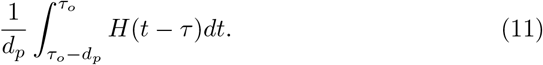

The age of any forager can be written in terms of the age of the youngest foragers. If we assume that the youngest foragers are aged *y*, then the age of any other forager can be written as *y* + *s*, where 0 ≤ *s* ≤ *τ*_*o*_ − *τ*_*y*_. Hence, a forager aged *y* +*s* entered pupation *τ*_*y*_ +*s* days ago and the time that has elapsed since this forager left pupation is *τ*_*y*_ + *s* − *d*_*p*_ (since *a* = *τ*_*y*_ + *s* − *d*_*p*_). Hence, for each age group of current foragers we have that the average number of hive bees present during their entire pupation is,

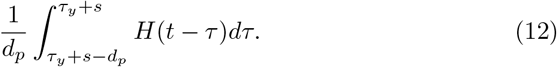

Finally, the average of the average hive bee populations over all forager ages is,

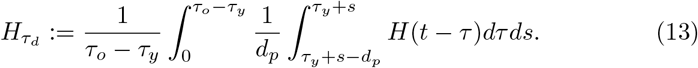

We use 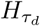 to track the average hive bee population present when the current foragers were capped brood and use 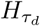 in the stress function. This function must have a maximum and a minimum value. The maximum corresponds to 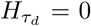 and the minimum occurs as 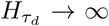. For large numbers of hive bees, where 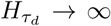, the hive has a sufficient number of hive bees to thermoregulate adequately and there should be no effect from stress. In this scenario the scaled components (of the model) should take their original values and so the minimum value of the stress function should be 1. Conversely, 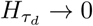 implies the hive has an insufficient number of hive bees to thermoregulate adequately and so the maximum effect of stress is felt. Let us denote this maximum effect of stress as *ψ*. We define the first stress function, which represents the impact of poor thermoregulation on current foragers, as,

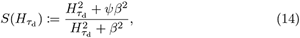

where *β* is the half saturation value, which is the number of hive bees required to experience half of the maximum effect of stress.

The second stress function can be derived directly from 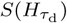. We noted earlier that the current hive bee population will have an increased food consumption rate as a result of actively thermoregulating, and so the second stress function is dependent on the current hive bee population. By replacing 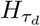 by *H* and *ρ*, respectively, in 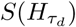 we define the current time stress function,

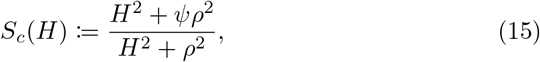

where *ρ* > *β* so that the curve saturates faster than 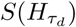. This assumption on *ρ* is due to a hives’ reaction to a need for thermoregulation. Initially, in colder temperatures, a hive will begin to experience a decrease in temperature and hive bees will begin to thermoregulate and so consume higher amounts of food. It is not till after the hive bees attempt to thermoregulate the hive that development of capped brood will be hindered as a result of poor thermoregulation. So, *S*_*c*_ saturates faster than *S*.

Now that we have constructed two stress functions 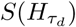 and *S*_*c*_(*H*) we can define the appropriate stress function for *S*_1_, *S*_2_ and *S*_3_. Since *S*_1_ and *S*_3_ model the impact of poor thermoregulation on foragers, due to suboptimal development as capped brood, we set 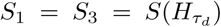. The function *S*_2_ models the increased food consumption of hive bees due to the need for thermoregulation. So we set *S*_2_ = *S*_*c*_(*H*). For the remainder of this paper we continue with the notation 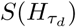 and *S*_*c*_(*H*).

### 2.3. Model with Thermoregulatory Stress

We have identified where in the Perry *et al.* model the effects of stress act and developed a function, 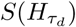, which represents the amount of stress as a result of poor thermoregulation during forager development. The three terms where thermoregulatory stress acts are; food collection rate,

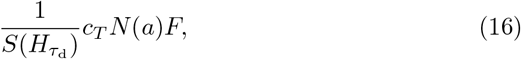

forager mortality rate,

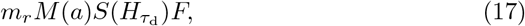

and the rate of hive bee food consumption,

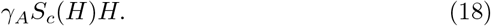

We incorporate these functions into the model of Perry *et al.* so that it becomes:

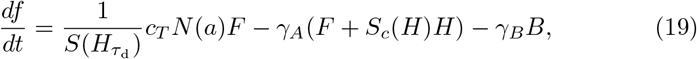

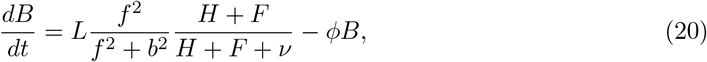

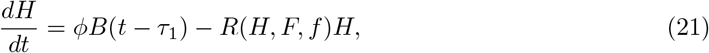

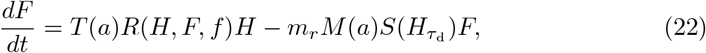

where *τ*_1_ is the total time spent in pupation. We have changed the dependence on hive bees of egg survival in the brood equation to depend on the entire adult bee population. With suitable adjustments of parameter values this is closely equivalent to using *H* only [22]. We also change the auxiliary functions: *N*(*a*), *T* (*a*) and *M*(*a*), so that all the functions are smooth and well defined,

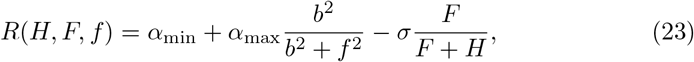

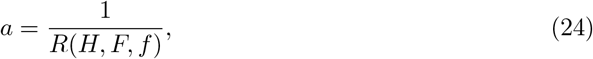

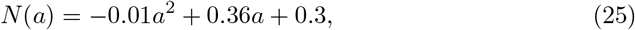

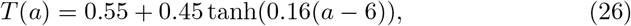

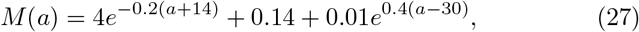

with *a* ∈ [2, 30]. We approximate the integral 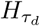 with a Reimann sum in numerical calculations,

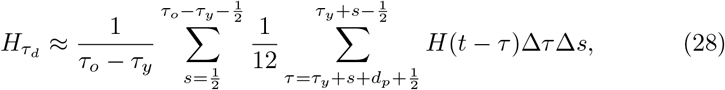

where ∆*τ* = ∆*s* = 1.

#### 2.3.1. Model Parameters

Many of the model parameters are taken from existing literature, particularly from [6]. We only need to determine parameter values for the new parameters introduced in the stress functions, 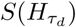 and *S*_*c*_(*H*).

The parameters which require values are *τ*_*y*_, *τ*_*o*_, *β*, *ρ*, *ψ* and *d*_*p*_. The delays *τ*_*y*_ and *τ*_*o*_ will change as the age of onset of foraging changes. However, for ease of calculation we will specify average values and use these in numerical solutions to (19)–(22). It was shown in [6] that worker bees are most efficient at foraging if they have reached 14 days of age as adults before starting to forage. We also know worker bees spend 12 days as capped brood. These two values suggest that approximately 26 days prior to *t*, the youngest foragers were brood entering pupation. So we set *τ*_*y*_ = 26 and *d*_*p*_ = 12. In order to determine *τ*_*o*_ we need to identify the average lifespan of a forager. That is the time from when an adult bee begins foraging till its death. This requires us to know the mortality of the current forager cohort. However we see later that this value itself is dependent on the delay, which creates a recursion. Hence, we approximate the average lifespan of a forager. Using results in [5, 6, 16] we conclude that a typical average mortality rate of a forager is 0.1 and so the average forager lifespan is 10 days. So we set *τ*_*o*_ = 36. Then

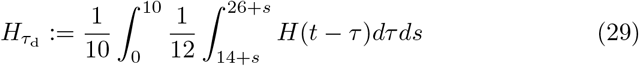

Finally we require values for *ψ*, *β* and *τ*. Perry *et al.* [6] suggested that the median rate of mortality for foragers in their model triples when the deleterious effects of precocious onset of foraging is included. We assume that the impact of sub-optimally developed bees is similar to that of precocious foraging and so we set *ψ* = 3. To determine *β* we identify the required number of hive bees to adequately thermoregulate the brood comb of a hive. Studies that count the number of hives bees distributed over the brood comb show that a ratio of one hive bee to approximately two capped brood cells is required for adequate thermoregulation [15]. Assuming that the average number of capped brood in a healthy hive is 16000, then the hive requires more than 8000 hive bees for adequate brood thermoregulation as hive bees cannot focus solely on thermoregulation. So, at approximately 8000 hive bees the hive will begin to experience thermoregulatory stress. We need a value of *β* so that the function *S*(*H*_*τ*_) approaches 1, for large *H*_*τ*_, slowly enough so that the effect of stress at *H*_*τ*_ = 8000 is slightly discernable. Hence, we choose *β* = 2000 (Figure 3 shows this behaviour). This choice of *β* leads us to setting *ρ* = 3000 in *S*_*c*_ in order to maintain the qualitative structure of the curve in Figure 3 but with faster saturation for *S*_*c*_(*H*) than for 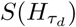.

**Figure 3:**
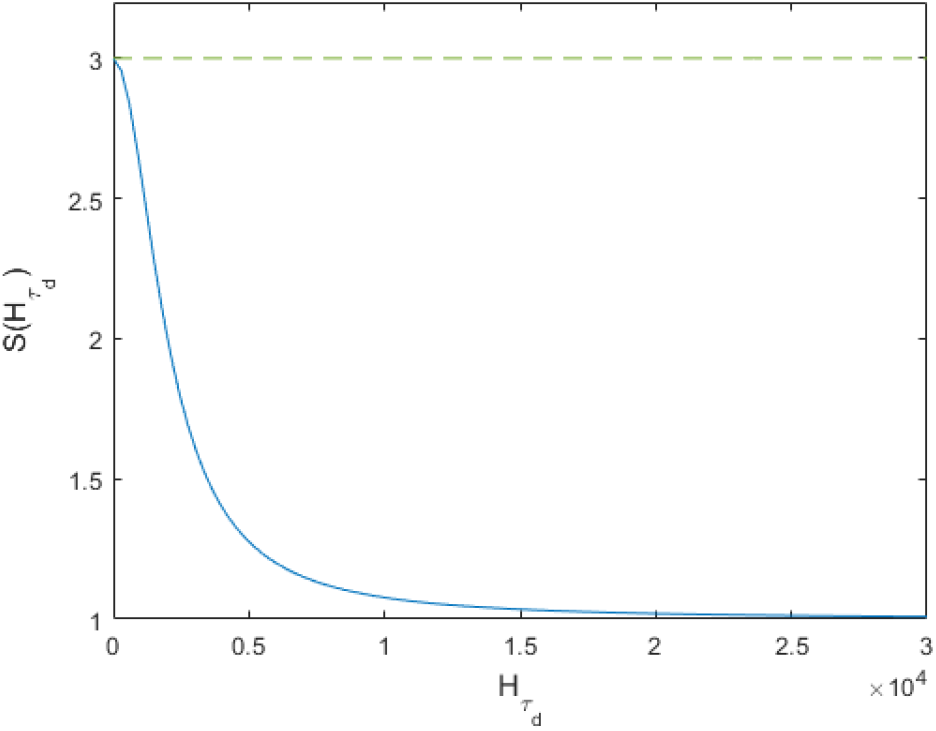
Stress functions 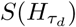 with *ψ* = 3 and *β* = 2000. The green dashed line indicates the maximum effect of stress.

## 3. Analysis and Results

To identify the existence of an Allee effect we must show a sensitivity to initial conditions in the system (19)–(22). The following analysis will show the existence of an Allee effect as well as showing that the existence is due to the inclusion of thermoregulatory stress in the model.

The system (19)–(22), is a set of closely coupled and nonlinear equations, and so analysis is difficult. In order to determine the qualitative behaviour of the system we generate a numerical bifurcation diagram (Figure 4). This diagram gives an indication of the stable equilibria for varying *m*_*r*_. By running simulations up to *t* = 6000 we are able to capture, numerically, the long time behaviour of the system. Figure 4 presents the bifurcation as the final value of the simulations, at *t* = 6000, when the system is assumed to have reached steady state, for different values of *m*_*r*_.

**Figure 4:**
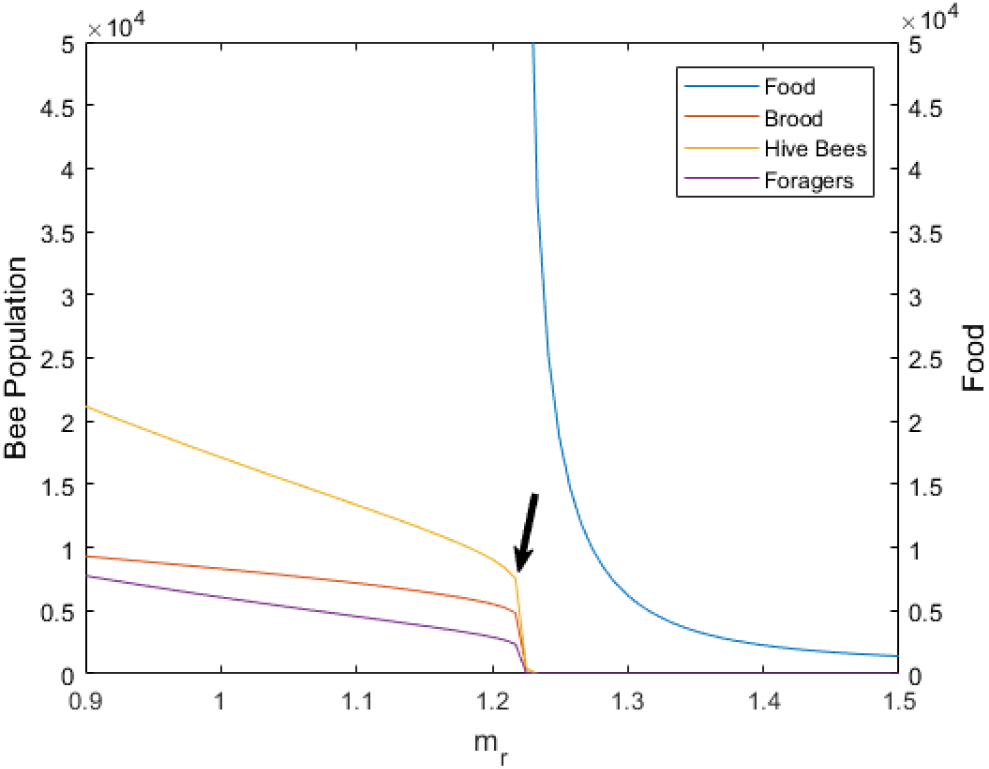
Numerical bifurcation diagram of the age-dependent system (19)–(22). All curves shown are stable equilibria. The black arrow indicates limit points in *B*, *H* and *F*.

We first analyse a reduced system by removing all terms in the system dependent on *a*. We then show that the analysis of this reduced system is a good approximation to the age-dependent system (19)–(22).

### 3.1. Analysis

Removing all age-dependent terms from (19)–(22) we obtain the following age-independent system,

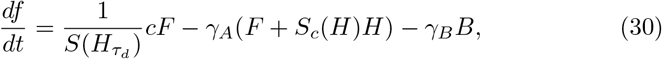

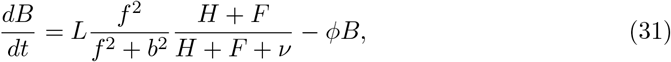

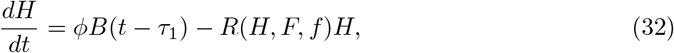

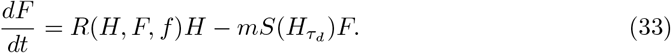

As well as removing the age-dependent functions we also change *c*_*T*_ to *c* and *m*_*r*_ to *m* as the former of both terms are ratios to scale food collection and mortality when expressed as a function of age. Here *c* is the amount of food collected per forager per day, and *m* is the forager mortality. Setting the LHS of equation (30) to zero gives,

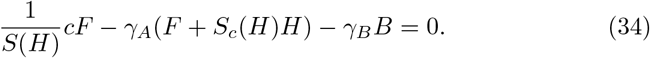

Rearranging (34) for *F* gives us an equation describing a surface,illustrated in Figure 5. This figure shows the values of *B*, *H* and *F* which satisfy (34) and hence give 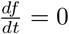. A trivial equilibrium solution occurs when *B*^∗^ = *H*^∗^ = *F*^∗^ = 0. We can show, numerically, that in fact the only equilibrium of (30)–(33), which satisfies (34), is *B*^∗^ = *H*^∗^ = *F*^∗^ = 0.

**Figure 5:**
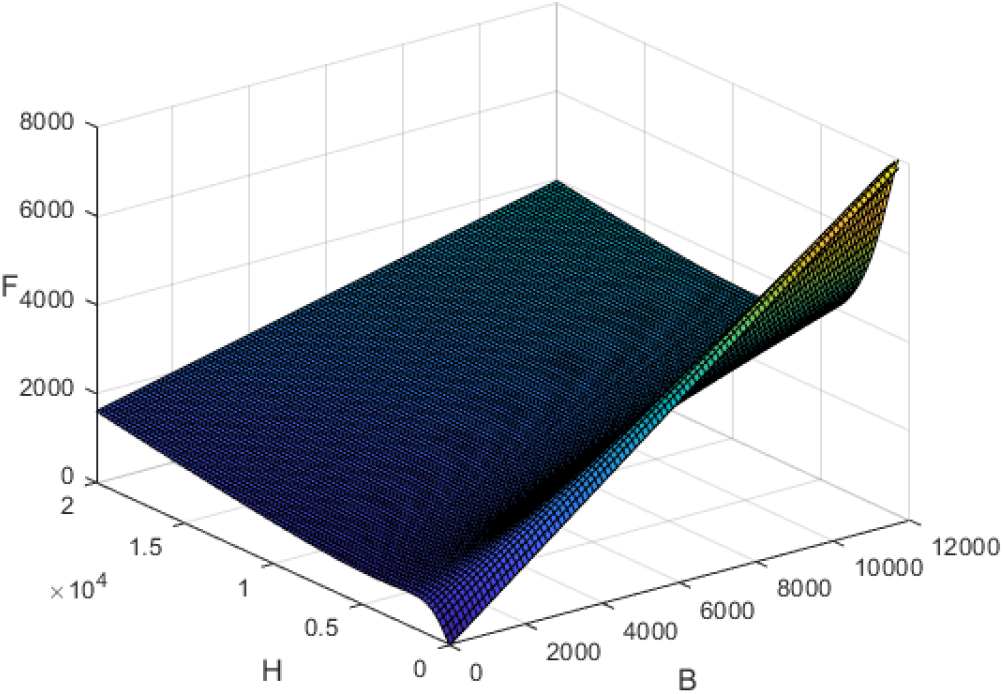
Surface in R^3^ which satisfies (34).

We let the LHS of (31) and (32) be zero and rearrange for *f*. We denote by *g*(*B, H, F*) the function derived from rearranging (31) and *k*(*B, H, F*) the function derived from rearranging (32) for *f*. We can show, numerically, that for each non-zero (in the sense that each element is non-zero) point (*B*_*s*_, *H*_*s*_, *F*_*s*_), where ⋅_*s*_ denotes a point on the surface (34), the value of *g*(*B, H, F*) ≠ *k*(*B, H, F*) for any (*B*_*s*_, *H*_*s*_, *F*_*s*_). Hence, the only coordinate of (34) that gives an equilibrium solution for (30)–(33) is (*B*_*s*_, *H*_*s*_, *F*_*s*_) = (0, 0, 0). However, *f* need not be zero. From (30)–(33), we see that for any non-zero *f*, all four equations will have zero as their RHS (as long as *B* = *H* = *F* = 0). This is consistent with the numerical bifurcation diagram, Figure 4, for the region to the right of limit point indicated in the diagram.

To determine what occurs in the system to the left of these limit points (indicated by the black arrow in Figure 4) we look at the limiting case in *f*. As suggested by the numerical bifurcation diagram, stored food tends to infinity as we approach the limit points from the right. This limiting approach was also used by Khoury *et al.* [16] to show stability of equilibria in their system. We take the limit *f* → ∞ in equations (31)–(33). This limiting case reduces the dimension of the system as equation (30) becomes decoupled from the system. It also reduces the complexity in the system as some terms simplify. In the limit as *f → ∞*, the system becomes,

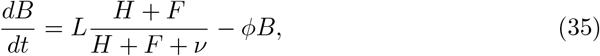

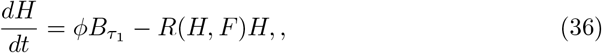

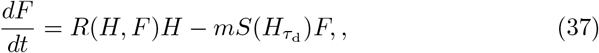

where 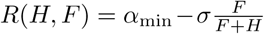. Equations (35)–(37) have up to three equilibria which can be found numerically. We use the infinitesimal generator numerical method [17] to find the largest eigenvalue of each steady state of system (35)– (37) for varying *m*. Figure 6 shows the largest eigenvalue for each equilibrium of the system with varying forager mortality in the range *m ∈* [0.1, 0.22]. Figure 6 shows the existence of three equilibria for 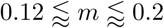. For *m* = 0.1 there is only one equilibrium. As *m* increases in the diagram, we see the creation of two additional equilibria via a saddle node bifurcation. One of these new equilibria eventually collides with the first equilibrium via another saddle node bifurcation at *m* ≈ 0.205 and we are left with one equilibrium once more. Figure 6 shows the value of the largest eigenvalue for each equilibrium, *λ*_max_.

In order to determine stability of the origin we first remove the singularity generated by the 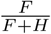 term in *R*(*H, F*). To do this we transform (35)–(37) to polar coordinates and use the infinitesimal generator approach to find the largest eigenvalue for the system in these coordinates. This calculation is laid out in detail in [19]. Figure 7 shows that the steady state at the origin changes stability at *m* ≈ 0.12.

**Figure 6:**
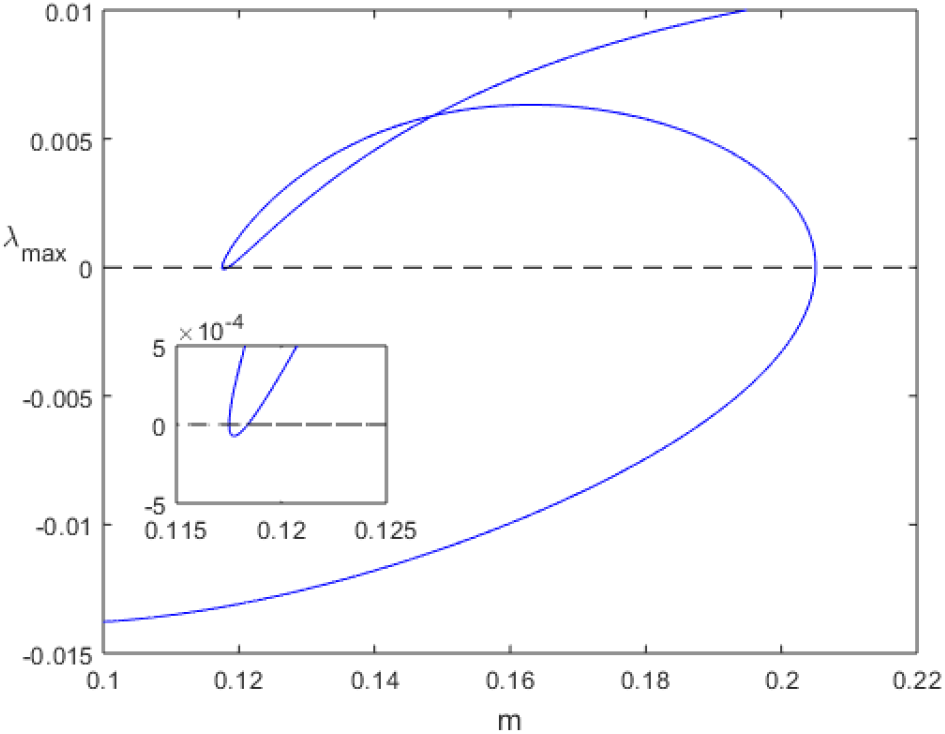
Largest eigenvalue of the infinitesimal generator for equilibria of system (35)–(37) as a function of *m*. When the largest eigenvalue is positive, the corresponding equilibrium is unstable, and when the largest eigenvalue is negative the corresponding equilibrium is stable. There are three equilibria when 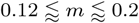.

**Figure 7:**
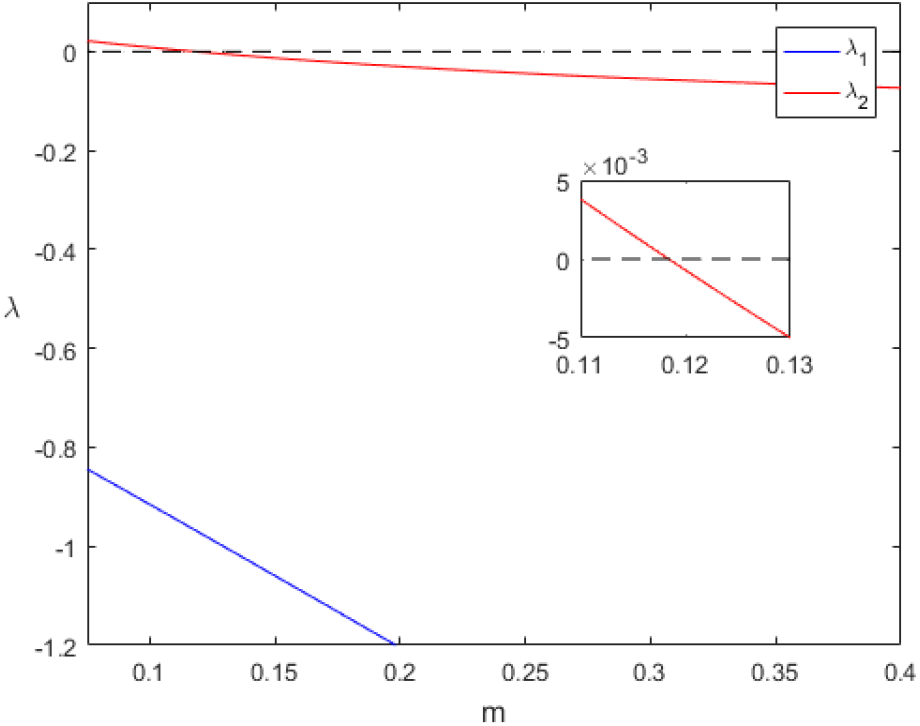
Largest eigenvalues, *λ*_1_ and *λ*_2_, of the infinitesimal generator for the origin of system (35)–(37).

We used the value of the largest eigenvalue of non-zero equilibria and the origin, as well as solving equations (35)–(37) numerically, to generate bifurcation diagrams for each variable. The bifurcation diagrams for each variable display the same qualitative behaviour so we choose *H*^∗^ as the measure of the solution. Figure 8 shows this diagram. There is a region of bistability between *m* ≈ 0.118 and *m* ≈ 0.205. Within this region the origin is a stable equilibrium and there is a second positive stable equilibrium. These are both separated by a third unstable equilibrium. Figure 8 shows that for any fixed *m* within this region there is an Allee effect and hence a sensitivity to initial conditions.

**Figure 8:**
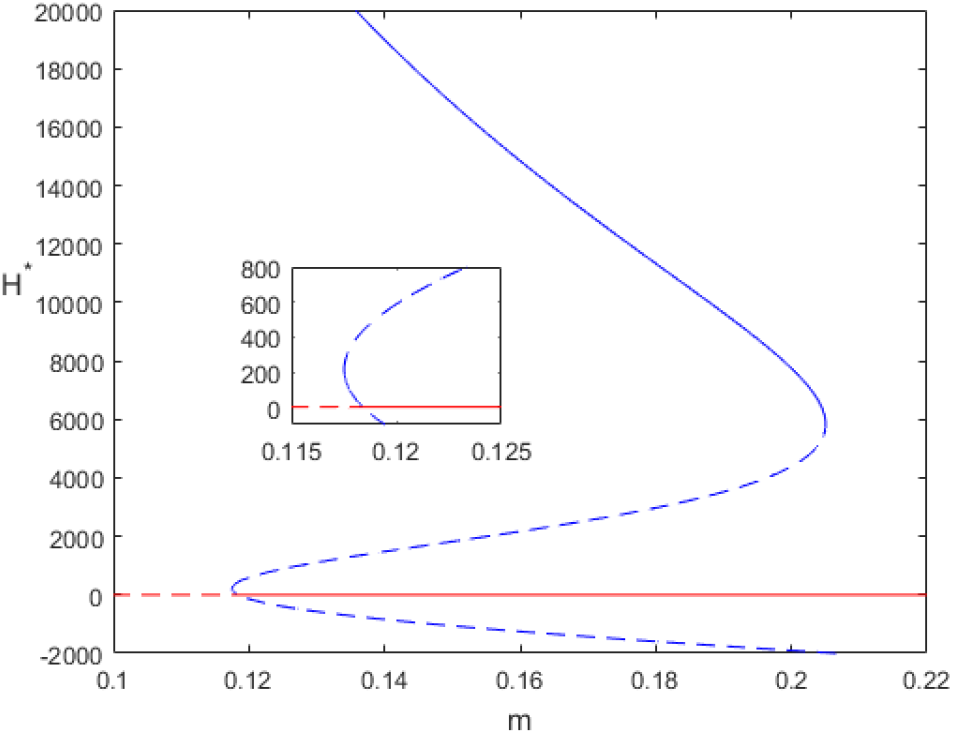
Bifurcation diagram for *H*^∗^ with parameter values as in Table 1. Dashed lines indicate unstable branches and solid lines indicate stable branches.

We can go further and show evidence that the source of this Allee effect in the age-independent model is due to the inclusion of thermoregulatory stress. By removing all age-dependent terms from (19)–(22) we recover the 4D model formulated by Khoury *et al.* [5], with the addition of stress due to poor thermoregulation. We have also made brood survival a function of *H* + *F* rather than *H* only as in [5]. Using the three population model (35)–(37) in the limit *f* → ∞, if we vary the values of *ψ* we recover the bifurcation diagram presented in [16] which corresponds to *ψ* = 1. In Figure 9 we see that at *ψ* = 1 the bifurcation in *H*^∗^ is qualitatively similar to that of the two population model presented in [16], with a transcritical bifurcation at the origin for *m* ≈ 0.35. When we set *ψ* > 1 the bifurcation curve has two folds and produces the bifurcation curve we see in Figure 8. This allows us to conclude that the source of the Allee effect in (30)–(33) is the inclusion of thermoregulatory stress.

**Table 1:** Parameter definitions and associated values for (19)–(22).

| Parameter | Meaning | Value |
| --- | --- | --- |
| $L$ | Queen's laying rate | 2000 |
| $\nu$ | Number of workers required in order for the number of new hive bees entering the hive from eclosion to be at half the rate of $L$ | 27000 |
| $\alpha_{\max}/\alpha_{\min}$ | Rate at which hive bees become foragers | 0.25 |
| $\sigma$ | Rate at which foragers revert to hive bees due to social inhibition | 0.75 |
| $\psi$ | Maximum effect on the death term from the effects of thermoregulatory stress | 2 |
| $\beta$ | Number of hive bees needed for the hive to experience half of the maximum effects of thermoregulatory stress | 2000 |
| $m_r$ | Ratio of forager mortality of an unhealthy hive to a healthy hive | 0.5-2 |
| $b$ | Half the amount of food needed for optimal egg survival | 500 |
| $c_T$ | Amount of food collected per forager per trip | 0.033 |
| $\phi$ | Rate at which brood enter pupation | 1/9 |
| $\gamma_A/\gamma_B$ | Food consumption per adult/brood respectively | 0.007/0.018 |
| $\tau_1$ | Time required for capped brood to emerge from pupation | 12 |

**Figure 9:**
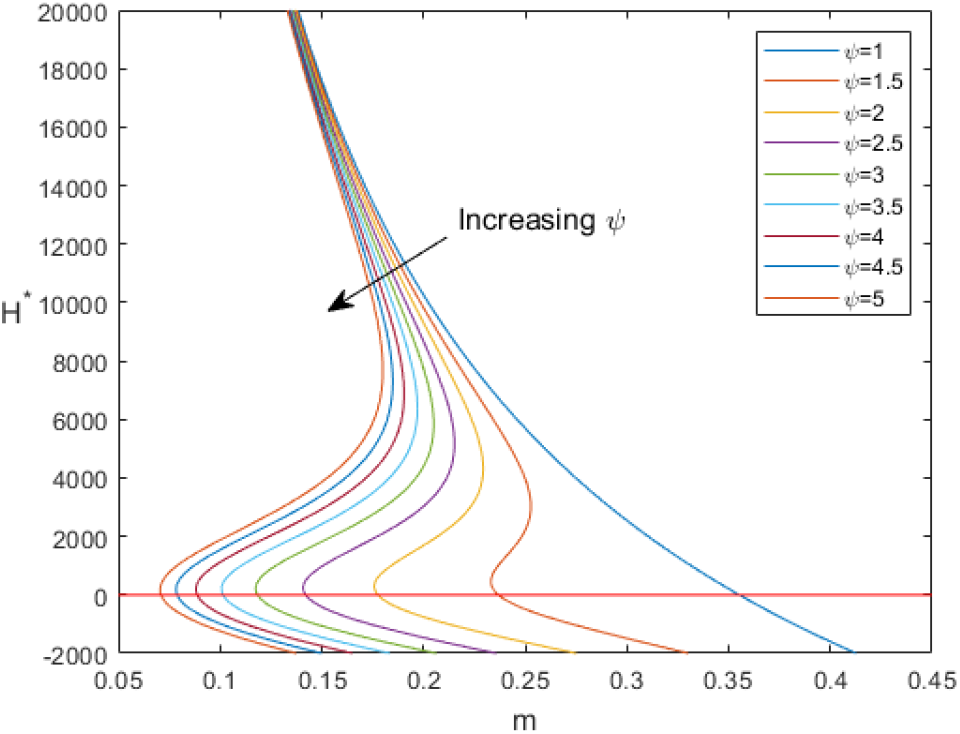
Changing bifurcation curve for varying values of *ψ*.

Finally, to determine how well the age-independent system (30)–(33) approximates the qualitative behaviours of the age-dependent system (19)–(22) we take the steady state values from Figure 4 and take the difference of each populations steady state values with the respective populations steady state values in (30)–(33). This difference is then normalised by dividing by the population size for each cohort in an average hive, for this we choose 15000 for brood, 45000 for hive bees and 15000 for foragers. Figure 10 shows that the normalised difference between the steady states of the two systems is always less that 10% and so we assume that the age-independent system (30)–(33) is a good approximation to the qualitative behaviour of the age-dependent system (19)–(22).

**Figure 10:**
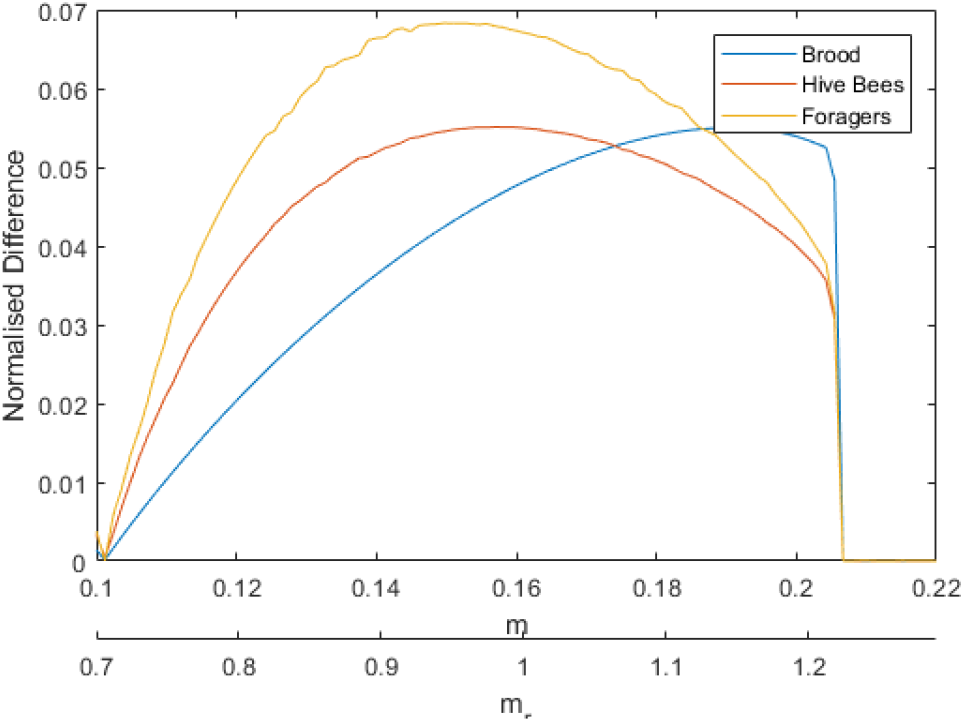
Normalised difference of approximation for bifurcation (absolute value of the difference of the numerical steady state values divided by the appropriate population value) diagrams between system (19)–(22) and system (30)–(33). We take the range [0.1, 0.22] for *m* and [0.7, 1.29] for *m*_*r*_. We use the values of 15000 brood, 45000 hive bees and 15000 foragers to normalise the error.

### 3.2. Simulations

The analysis of the age-independent system (30)–(33) suggests that the age-dependent system will display a sensitivity to initial conditions via an Allee effect. We can show the existence of this Allee effect by solving the age-dependent system numerically and exploring the way that outcomes of the model depend on the initial hive population. Figure 11 shows that for two different initial sets of hive populations, with fixed *m*_*r*_, the hive either survives or collapses depending on the initial condition. This confirms the existence of an Allee effect.

**Figure 11:**
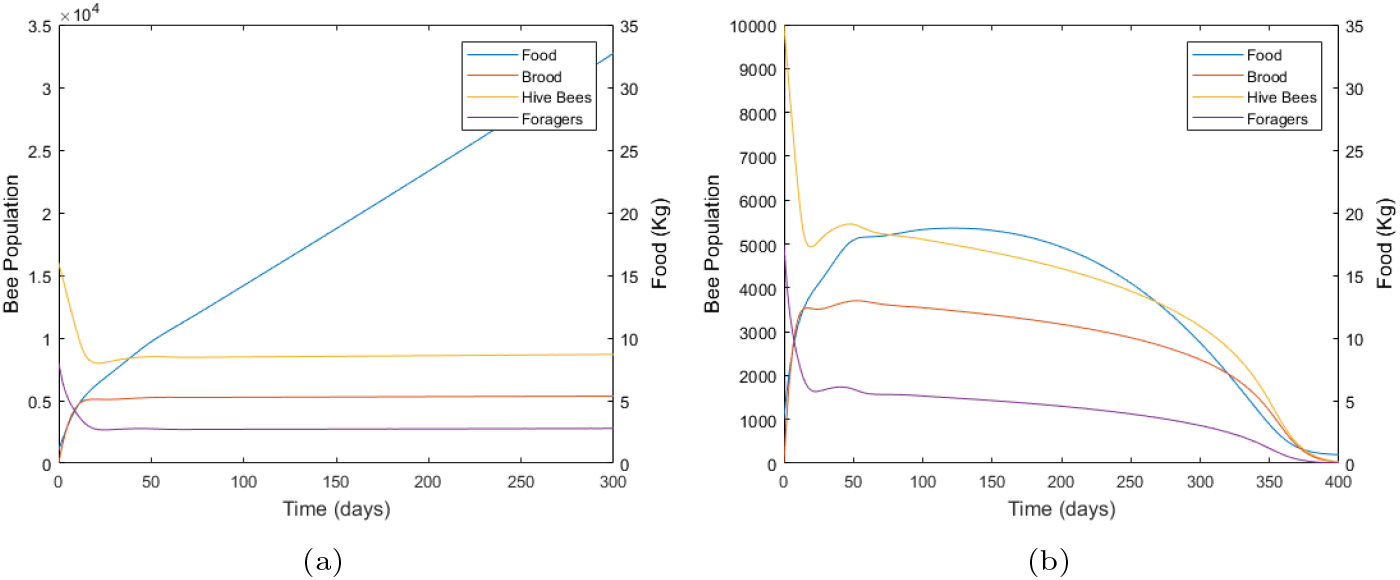
Numerical solutions for hive populations as a function of time that demonstrates an Allee effect for initial condition (a) (*f*_0_, *B*_0_, *H*_0_, *F*_0_) = (1000, 0, 16000, 8000) and (b) (*f*_0_, *B*_0_, *H*_0_, *F*_0_) = (1000, 0, 10000, 5000). In both simulations *m*_*r*_ = 1.2 and parameter values used are as in Table 1.

Another outcome of the analysis of system (30)–(33) was that the final populations in the hive is independent on the amount of food stored in the hive. We use simulations to provide evidence of the independence of the final state of the system from the amount of stored food. In Figure 12 we manipulate the amount of food at *t* = 150 days by halving the amount of food in the surviving hive and doubling the amount of food in the collapsing hive from simulations in Figure 11. Figure 12 shows that this has no effect on the outcome for either hive. The surviving hive continues to prosper and the collapsing hive ultimately collapses. This also shows that the collapse is not driven by stored food availability.

**Figure 12:**
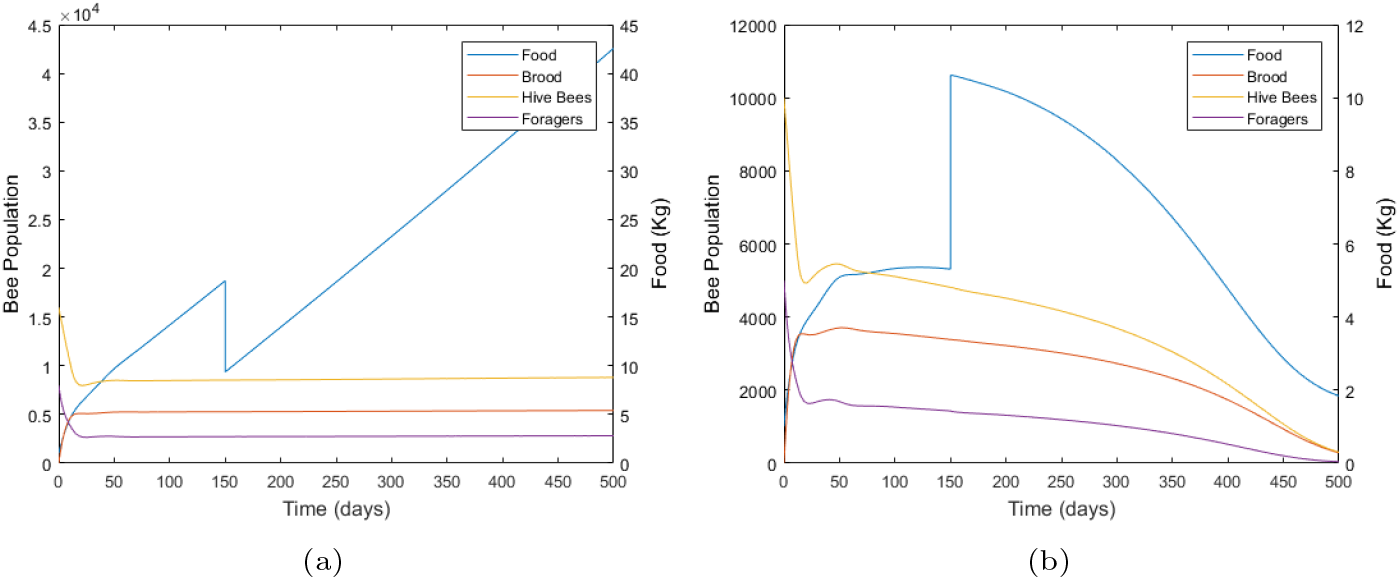
Numerical solutions for initial condition (a) (*f*_0_, *B*_0_, *H*_0_, *F*_0_) = (1000, 0, 16000, 8000) and (b) (*f*_0_, *B*_0_, *H*_0_, *F*_0_) = (1000, 0, 10000, 5000) with *m*_*r*_ = 1.2 and parameter values as in Table 1 in both simulations. The amount of food is halved and doubled in (a) and (b) respectively. The variation in the amount of food in each simulation does not alter the outcome of the hive.

The choice in the amount of change in stored food for simulations in Figure 12 is arbitrary. We chose an amount which would be significant in a real world setting. There is an upper limit to the amount of food that can be removed from a hive, in our simulations, for which the outcome of the hive does not change. Comparing the simulations in Figure 12(a) and Figure 13(a), the ratio of food removed to stored food prior to removal is larger in the latter yet there is no change in the qualitative outcome of the hive. So this upper limit on the amount of food that can be removed is so large that we can assume that it is irrelevant for practical purposes. Likewise, by choosing an arbitrarily large increase in the amount of food (Figure 13(b)) the hive will continue to collapse. Therefore, we can assume that in a real world scenario increasing or decreasing the amount of food for a collapsing or sustained hive, respectively, will have no outcome on the qualitative health of the hive long term if thermoregulatory stress is present.

**Figure 13:**
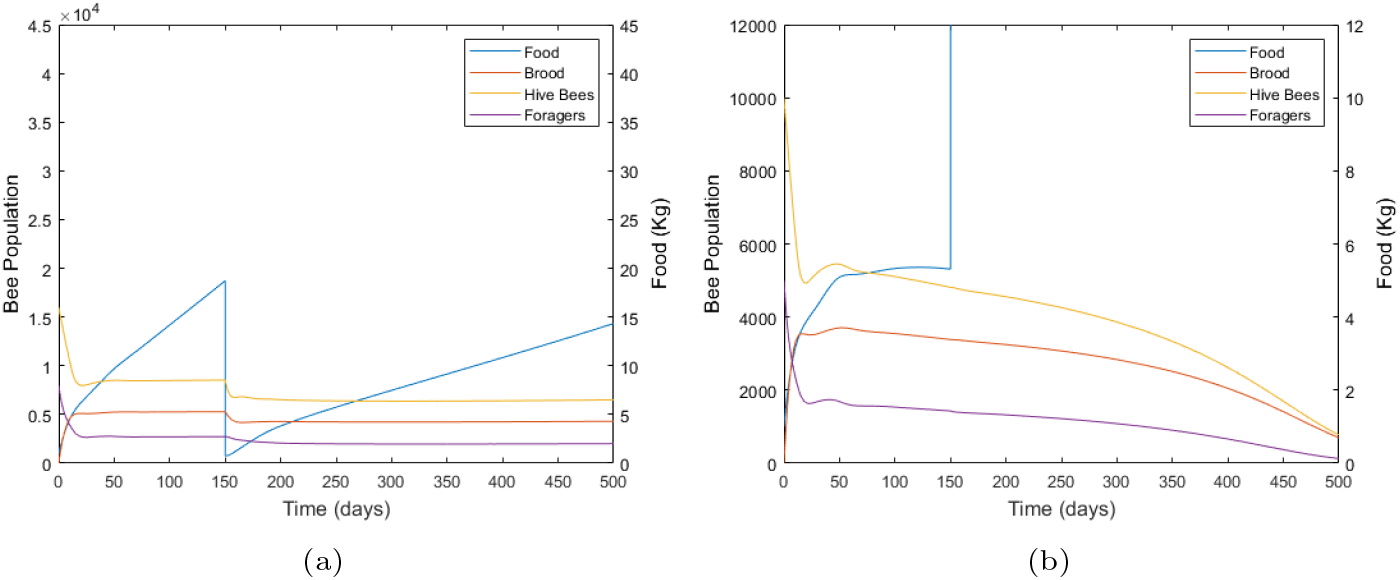
Numerical solutions for initial condition (a) (*f*_0_, *B*_0_, *H*_0_, *F*_0_) = (1000, 0, 16000, 8000) and (b) (*f*_0_, *B*_0_, *H*_0_, *F*_0_) = (1000, 0, 10000, 5000) with *m*_*r*_ = 1.2 and parameter values as in Table 1 in both simulations. The amount of food is reduced by 96% in (a) and increased by 200 times its amount in (b).

The model with thermoregulatory stress predicts that the inclusion of stress generates an Allee effect. We also see that while manipulating the amount of stored food in a hive does not have any major impact on the qualitative health of the hive, it does extend the time to collapse in an already collapsing hive.

## 4. Discussion

We have developed a model that specifically represents the stress due to a breakdown in thermoregulation. This new model is based on the model in [6] but is focused on the effects of temperature stress. By analysing the equations we showed that in this model thermoregulatory stress produces an Allee effect (Figure 9). Two saddle-node bifurcations are created as the model changes from the case where there is no stress (*ψ* = 1) to where thermoregulatory stress acts on the hive (*ψ* > 1). This Allee effect can be observed in simulations (Figure 11) where there are different outcomes for two different initial conditions applied to the same model with the same parameter set.

There are several models which include stress in a hive [8, 7]. These models incorporate stress by using per capita death rates or simply adding a death term to a population equation. Honey bees have been well studied and documented for many years. This allows us to explicitly model sources of stress. A break-down in thermoregulation occurs as a result of low hive bee population, and is known to negatively impact hive health and dynamics. Poor thermoregulation of capped brood leads to under-developed workers which are not as capable at foraging as their optimally developed counterparts. For hive bees to be able to keep the hive at optimal temperature, particularly in cold weather, they require more food as the task increases the bees’ energy requirements and outputs. To model these biological processes we created a model which includes the effects of poor thermoregulation on forager mortality and on food stores. This is achieved by identifying how past behaviour of hive bees affects the current population of foragers in the hive and, in particular, how the hive’s ability to thermoregulate in the past, when the current forager were capped brood, determines the efficacy of the current foragers. We also modelled the current time food consumption of hive bees as they increase their metabolism in order to regulate the internal temperature of the hive. The inclusion of models for these specific stresses produces a faster decline in populations than for the model with no thermoregulatory stress.

By modelling stress explicitly in (19)–(22), we can make connections between simulations of model hives and data recorded from real hives experiencing thermoregulatory stress. This provides an immediate and clear advantage over models that include stress by adding generic terms, such as those in [7, 8]. We also observe that modelling stress explicity in our model produces an Allee effect similar to that seen in [8]. We can potentially use our model, not only to determine how a hive currently experiencing thermoregulatory stress may behave in the future, but also to examine what actions can be taken to rescue a hive, that is failing due to a breakdown in thermoregulation. The explicit inclusion of stress means the model can be analysed to examine how different castes in a hive are affected by the stress and how the stress may be reduced or amplified by bees’ behaviour.

The analysis of (19)–(22) showed that the model hives’ dependence on food is less crucial than the effect of population demography. Figure 5 shows, numerically, that the only steady state of the system is when *B*^∗^ = *H*^∗^ = *F*^∗^ = 0. However, when *f* → ∞ then *B*, *H* and *F* all reach a steady state value. This suggests a multiple time scale dynamic [19] and that food limitation does not drive population numbers, at least in this model. This independence of survival from food is observed, for example, in rental hives used in pollinating crops [19, 23]. These small hives (1.4-2.7Kg [23]) which are placed in multiple locations through plantations to pollinate crops suffer significant losses. Even when apiarists supply large amounts of pollen and sucrose solutions to the hives before they are deployed, the hive still suffers significant losses of their adult bee populations albeit not as severe as without the additional pollen and sucrose solution. This is similar to the final outcome of simulations in Figure 11(b) and 12(b) where initial colonies are both the same, and the addition of food 150 days into the simulations in Figure 12(b) merely serves to delay the same outcome of hive loss.

The lack of food availability drives the rapid decline in hive populations in the models of [5, 6]. The explicit inclusion of stress in our model shows that food is no longer the major driver of collapse, although, we see much of the same symptoms associated with collapse in both models. The rapid decline in age at onset of foraging (AAOF) in [6] is also observed in our results. Figure 14(a) shows the change in AAOF with varying *m*_*r*_. We see, as expected, that the decline in AAOF in Figure 14(a) occurs for the same value of *m*_*r*_ as the increase in the amount of stress seen in Figure 14(b). In [6] the authors examined the median AAOF as a function of *m*_*r*_. The authors show that there is a significant crash in the median AAOF for *m*_*r*_ ≈ 1.9 where AAOF rapidly declines from approximately 9 days to the minimum value of 2 days. Equations (19)–(22) include the effects of thermoregulatory stress where we know this acts in a hive. By examining AAOF in (19)–(22) we notice a sharp decline at *m*_*r*_ ≈ 1.25 where AAOF rapidly declines from 14 days to 6 days. While the system presented in this paper exhibits a similar declining behaviour as that in [6] the range of AAOF where the decline occurs is slightly greater here than that in [6] and the decline of AAOF in our model occurs much earlier.

**Figure 14:**
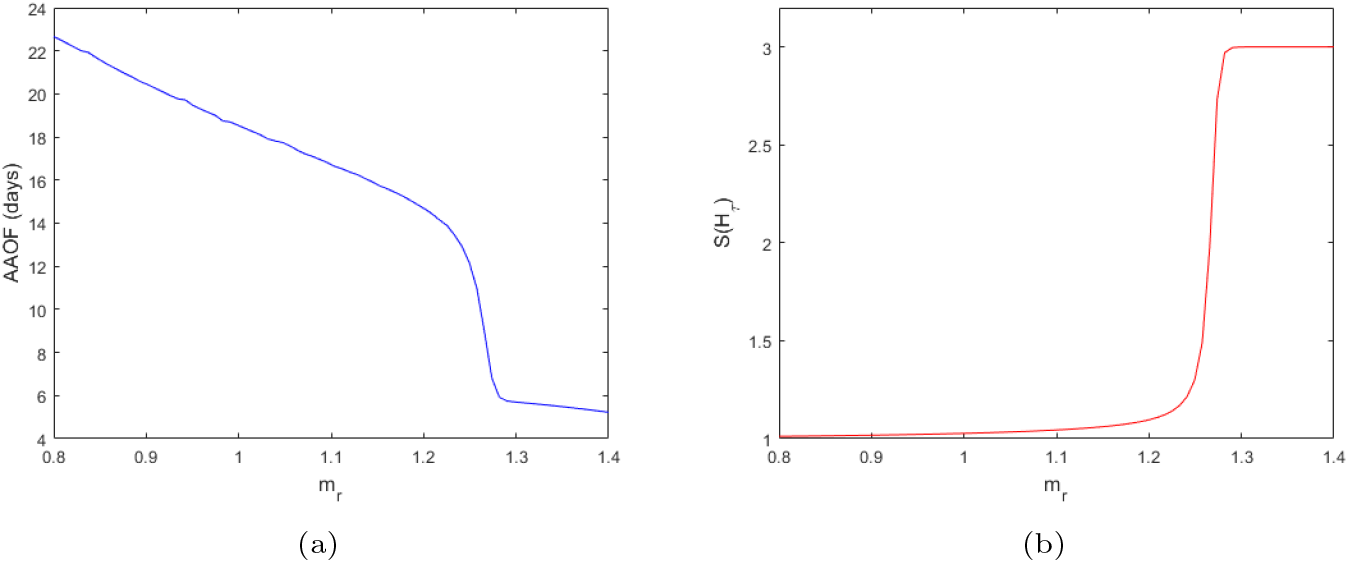
Steady state value of (a) age at onset of foraging and (b) thermoregulatory stress for varying *m*_*r*_. Initial condition uses is (*f*_0_, *B*_0_, *H*_0_, *F*_0_) = (1000, 0, 16000, 8000) with parameter values as in Table 1. In both plots, simulations were allowed to run for 600 days.

The model presented in this paper is a first step in explicitly modelling stressors on a honey bee hive. We showed that the inclusion of thermoregulatory stress produces an Allee effect. The model also predicts that ultimately the demography of a hive outweighs other factors in a hives’ ability to survive, although there are other factors which may temporally prolong a hive’s survival, including food availability inside the hive. The idea of this paper can be extended by including other known stressors which reduce the efficacy of bees. If worker bees cannot perform their tasks well, the hive as a whole suffers which eventually leads to significant losses, if not a collapse. The inclusion of thermoregulatory stress, explicitly, gives us new insight into what may be happening inside the hives themselves. The explicit inclusion of thermoregulation allows us to identify where and why collapses occur in model hives which gives direct insight into what may occur in real hives.

## References

[1] Vanengelsdorp D,Evans J.D, Saegerman C, et al., 2009, Colony collapse disorder: A descriptive study. PloS ONE, 4(8), e6481.

[2] Oldroyd B.P, 2007, What’s killing American honey bees? PLoS Biology, 5(6), e168.

[3] Ratnieks F.L.W, Carrack N.L, 2010, Clarity on honey bee collapse? Science, 327(5962), pp. 152–153.

[4] Goodwin R.M, Cox H.M, Taylor M.A, et al., 2011, Number of honey bee visits required to fully pollinate white clover (*Trifolium repens*) seed crops in Canterbury, New Zealend. New Zealand Journal of Crop and Horticultural Science, 39(1), 7–19.

[5] Khoury D.S, Barron A.B, Myerscough M.R, 2013, Modelling food and population dynamics in honey bee colonies. PLoS ONE, 8(5), e59084.

[6] Perry C.J, Sovik E, Myerscough M.R, Barron A.B, 2015, Rapid behavioral maturation accelerates failure of stressed honey bee colonies. PNAS, 112(11), 3427–3432.

[7] Bryden J, Gill R.J, Mitton R.A.A, Raine, N.E, Jansen V.A.A, 2013, Chronic sublethal stress causes bee colony failure. Ecology Letters, 16, 1463–1469.

[8] Booton R.D, Iwasa Y, Marshall J.A.R, Childs D.Z, 2017, Stress-mediated Allee effects can cause the sudden collapse of honey bee colonies. Journal of Theoretical Biology, 420, 213–219.

[9] Betti M.I, Wahl L.M, Zamir M, 2014, Effects of Infection on Honey Bee Population Dynamics: A Model. PLoS ONE, 9(10), e110237.

[10] Kronenberg F, Heller H.C, 1982, Colonial thermoregulation in honey bees (*Apis mellifera*). Journal of Comparative Physiology, 148(1), 65–76.

[11] Jones J.C, Helliwell P, Beekman M, et al., 2005, The effects of rearing temperature on developmental stability and learning and memory in the honey bee, *Apis mellifera*. Journal of Comparative Physiology A, 191(12), 1121–1129.

[12] Stabentheiner A, Kovac H, 2016, Honeybee economics: optimisation of foraging in a variable world. Scientific Reports, 6(28339).

[13] McMullan J.B, Brown M.J.F, 2005, Brood pupation temperature affects the susceptibility of honeybees (*Apis mellifera*) to infestation by tracheal mites (*Acarapis woodi*). Apidologie, 36(1), 97–105.

[14] Tautz J, Maier S, Groh C, Rossler W, Brockmann A, 2003, Behavioral performance in adult honey bees is influenced by the temperature experienced during their pupal development. PNAS, 100(12), 7343–7347.

[15] Delaplane K.S, van der Steen J, Guzman-Novoa E, 2013, Standard methods ofr estimating strength parameters of *Apis mellifera* colonies. Journal of Apicultural Research, 52(1).

[16] Khoury D.S, Myerscough M.R, Barron A.B, 2011, A Quantitative Model of Honey Bee Colony Population Dynamics. PLoS ONE, 6(4), e18491.

[17] Breda D, Maset S, Vermiglio R, 2015, Stability of Linear Delay Differential Equations A Numerical Approach with MATLAB. Springer New York.

[18] Kulhanek et al.,2017, A National Survey of Managed Honey Bee 2015-2016 Annual Colony Losses in the USA. Journal of Apicultural Research, 56(4), 328–240.

[19] Zeaiter Z, 2019, The Effect of Thermoregulatory Stress on Honey Bee Hives (Unpublished masters thesis). The University of Sydney, Sydney, Australia.

[20] Colin T, Meikle W.G, Paten A.M, Barron A.B, 2019, Long-term dynamics of honey bee colonies following exposure to chemical stress. Science of The Total Environment, 677, 660–670.

[21] Colin T, Meikle W.G, Wu X, Barron A.B, 2019, Traces of a Neonicotinoid Induce Precocious Foraging and Reduce Foraging Performance in Honey Bees. Environment Science & Technology, 53(14), 8252–8261.

[22] Khoury D.S, 2009, Colony collapse disorder: Quantitative models of honey bee population dynamics (Unpublished honours thesis). The University of Sydney, Sydney, Australia.

[23] The University of Maine, 2013. Honey Bees and Blueberry Pollination, 629.

